# Landscape of RNA editing reveals new insights into the dynamic gene regulation of spermatogenesis based on integrated RNA-Seq

**DOI:** 10.1101/478206

**Authors:** Xiaodan Wang, Zhenshuo Zhu, Xiaolong Wu, Hao Li, Tongtong Li, Qun Li, Peng Zhang, Leijie Li, Dongxue Che, Xia Xiao, Jinlian Hua, Mingzhi Liao

## Abstract

Spermatogenesis is an important physiological process associated with male infertility. But whether there are RNA editings (REs) and what’s the role of REs during the process are still unclear. In this study, we integrated published RNA-Seq datasets and established a landscape of REs during the development of mouse spermatogenesis. 7530 editing sites among all types of male germ cells were found, which enrich on some regions of chromosome, including chromosome 17 and both ends of chromosome Y. Totally, REs occur in 2012 genes during spermatogenesis, more than half of which harbor at two different sites of the same gene at least. We also found REs mainly occur in introns, coding regions (CDSs) and intergenic regions. Moreover, about half of the REs in CDSs can cause amino acids changes. Finally, based on our adult male Kunming mice, we verified that there is a non-synonymous A-to-I RNA editing site in *Cog3* during spermatogenesis, which is conserved not only between species but also across tissues. In short, based on the power of integrating RNA-Seq datasets, we provided the landscape of REs and found their dynamic changes during mouse spermatogenesis. This research strategy is general for other types of sequencing datasets and biological problems.

## INTRODUCTION

Infertility is gaining increasing attention which affects about 15% couples all over the world, among which male factor is responsible, alone or in combination with female factors in about half of the cases(Agarwal et al., 2015; JP et al., 2002; Pizzol et al., 2014). Dysfunction of sperms may arise from different factors, including lifestyle, heredity, obesity, drugs, and so on. Among them, genetic abnormalities account for 15%–30% of male infertility(Ferlin, 2012; O'Brien et al., 2010). Genetics leads to male infertility by influencing many physiological processes, such as spermatogenesis, sperm quality and hormonal homeostasis. Therefore, revealing the molecular mechanisms of spermatogenesis is essential for the understanding of male infertility. Spermatogenesis is a complex and dynamic process leading to the continuous production of sperm(Che et al., 2017). This process requires the successive and coordinated expression of thousands of genes, mixed with multi-level regulations from transcriptional, post-transcriptional and translational gene regulation(Bai et al., 2017).

As a kind of post-transcriptional regulation, RNA editings (REs) change the genetic information at the mRNA level, providing flexible regulation model from mRNA to proteins and making organisms to respond environment changes quickly and easily. Transcripts of some genes should be edited to initiate an effective translation according to the need of specific tissues and developmental times. Most of the previous studies on REs focused on A-to-I editing, which is most common in humans and other primates and is often associated with cancer(Paz et al., 2007; Slotkin and Nishikura, 2013; Wang et al., 2017).Generally, REs occur more frequently in non-coding than in coding regions. In human, RNA editing mainly occurs in introns and 5’- or 3’-untranslated regions (UTRs)(Athanasiadis et al., 2004; Blow et al., 2004; Kim et al., 2004). Considering the complex and dynamics of spermatogenesis and the flexible regulation manner of REs, we supposed there may be close relationships between them. Though some hints have showed REs play a certain roles during spermatogenesis, no evidence supports the existence of REs, let alone the global perspective of REs function in germ cells.

To address the above problems, we integrated all published RNA-Seq datasets and calling REs related with different types of cells during spermatogenesis. Interesting, we found there are huge amount of REs in specific cells and the dominant types of REs are not only A-to-I, but also G-to-A, C-to-U and U-to-C editing in male germ cell process of mouse. Different with human, we found REs mainly occur in introns, CDSs and intergenic regions in mouse. Even more, about half of the editing events in CDSs result in amino acid changes. We also found several novel genes with non-synonymous REs are related with different male germ cells, including *Rnf17*, *Boll*, *Adad1* and *Rbmy*. As a typical example, we verified non-synonymous REs in *Cog3,* which is conserved not only between species but also across tissues. In summary, based on our integrated analysis and landscape of REs, we found that REs play key roles during spermatogenesis.

## RESULTS

### Data preprocessing

The framework of our works was shown as Fig. 1. We collected published 12 studies of RNA-Seq datasets about spermatogenesis from Gene Expression Omnibus (GEO), Sequence Read Archive (SRA). These datasets covered 100 samples and were all sequenced with illumine machines platform. In addition to the previous 100 samples, we also collected the latest published datasets under accession number GSE75826 as validation data. Our integration strategy has increased the depth of sequencing to some extent, while at the same time obtaining consistent editing sites from different samples. Compared to detecting RNA editing sites in each sample separately, this method may improve accuracy. After data preprocessing, we identified the nucleotide changes and removed sites with overlap regions of positive and negative chains. Finally, we eliminated single-nucleotide polymorphisms (SNPs) and determined the final list of REs sites. Comparing the results of the integrated and non-integrated samples under the same identify process, we found that fewer editing sites identified from integrated samples than the unintegrated samples, but the editing sites identified by the integration method were consistent across samples. What’s more, some editing sites cannot be detected in single samples due to the low sample sequencing depth, but can be identified in the integrated samples. Calculate the Simpson's diversity index of integration and non-integration results, with a minimum of 0.65 and a maximum of 0.98(Supplemental Fig. S1).

**Figure 1.**
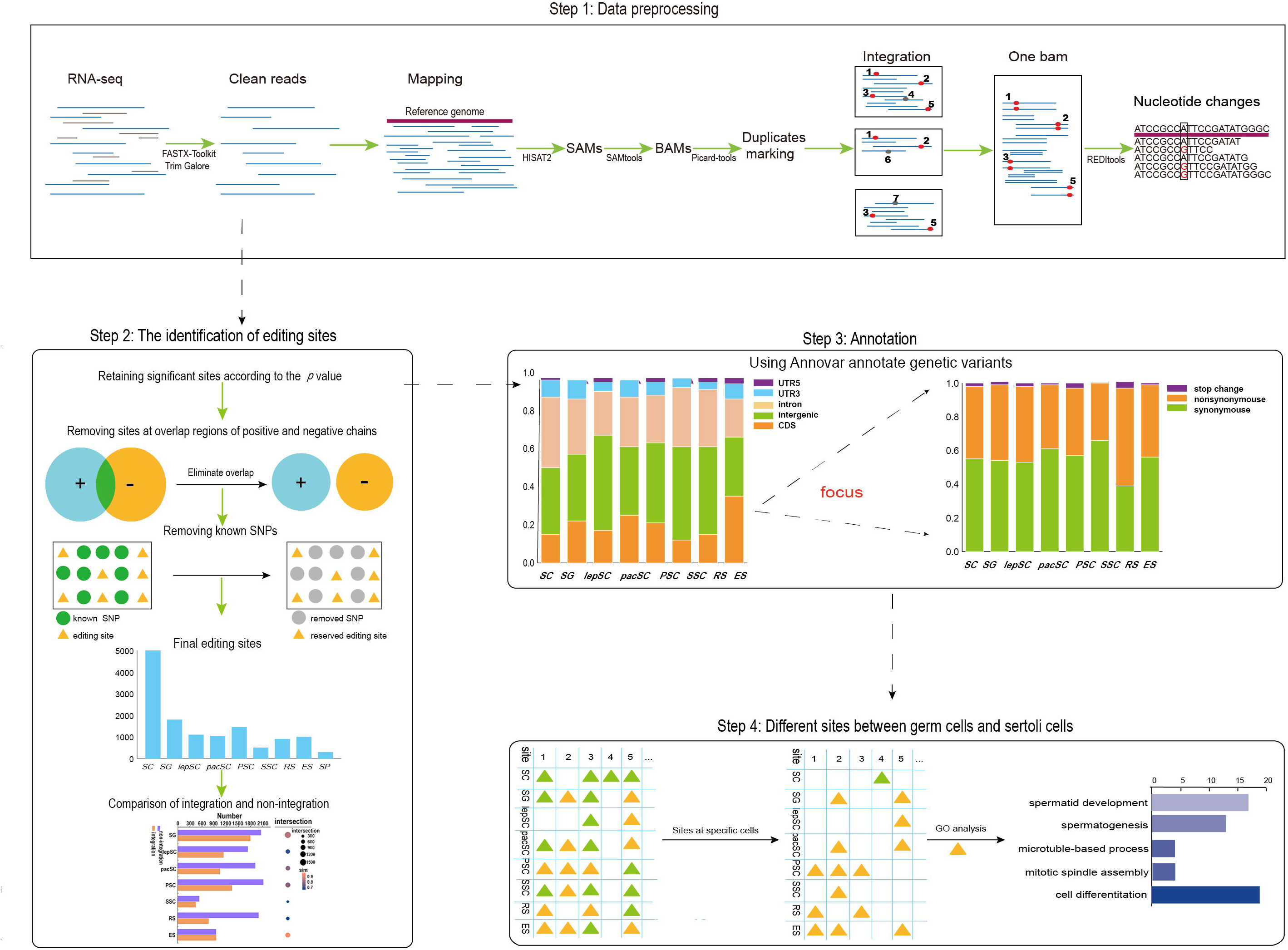
Experiment process. The whole experiment can be divided into four parts: data preprocessing, the identification of editing sites, genetic variants annotation and getting cell-specific editing sites.

### Identifying editing sites in specific cells during spermatogenesis

Based on the developmental stages, cells in testis were divided into nine different kinds(8 kinds of male germ cells and sertoli cells as control) (Supplemental Table S1), including spermatogonia (SG), leptotene spermatocytes (lepSC), pachytene spermatocytes (pacSC), primary spermatocytes before meiosis LJ (PSC), secondary spermatocytes (SSC), round spermatids (RS), elongative spermatids (ES) and sperm (SP). Results showed that the number of REs in sertoli cells (SC) is significantly larger than in germ cells (Fig.2A; Table 1). It may be due to the activation of RNA synthesis in sertoli cells. On the other hand, the meiosis process germ cells will present chromatin condensation, diploid into haploid, nuclear degeneration, which will affect the gene expression, reduce the amount of RNA and even terminate RNA synthesis. When the sperm cells begin to elongate, RNA synthesis gradually ceases, and there is no RNA synthesis in mature spermatozoa. A considerable proportion of the RNA synthesized during the pachytene stage is preserved through spermatid development until late spermiogenesis(Geremia et al., 1977; Monesi, 1965; Moore, 1971). Therefore, we did not analyze the RNA editing sites identified in mature spermatozoa in the subsequent analysis. Finally, 7530 editing sites among other 7 types of male germ cells were found (Fig.2A; Table 1). According to the editing level distribution of the editing sites in each cell, male germ cells have higher RNA editing level than sertoli cells(Fig.2B;Fig.2C). By the way, though the samples’ numbers of different kinds of cells were different, there was not significant association with the number of editing sites (Fig. 2D), which indicates the robust results among different scales of sample size. We divided all editing sites into seven major modules based on the type of cells they exist (Fig. 2E). The results showed that many editing sites were cell-specific, such as the performance of editing events on *Nudt6*, *Chd5*, *Wdfy1*, *LOC101056073*, *Foxn3*, *Cnot10* and *Zfp534* genes (Fig. 2E). It is indicated that many RNA editing events occur at specific stages of spermatogenesis and are dynamic, which is consistent with the dynamic expression of genes during spermatogenesis.

**Table 1.** The details of the editing sites identified in specific cells

|  | SC |  | SG |  | lepSC |  | pacSC |  | PSC |  | SSC |  | RS |  | ES |  | SP |  |
| --- | --- | --- | --- | --- | --- | --- | --- | --- | --- | --- | --- | --- | --- | --- | --- | --- | --- | --- |
|  | Editin<br>g<br>sites | #gene<br>s | Editin<br>g sites | #gene<br>s | Editin<br>g sites | #gene<br>s | Editin<br>g sites | #gene<br>s | Editin<br>g sites | #gene<br>s | Editin<br>g sites | #gene<br>s | Editin<br>g sites | #gene<br>s | Editin<br>g sites | #gene<br>s | Editin<br>g sites | #gene<br>s |
| All regions | 5402 | 2008 | 1803 | 897 | 1140 | 621 | 1047 | 650 | 1349 | 700 | 455 | 364 | 775 | 521 | 961 | 478 | 304 | 190 |
| Exonic | 796 | 316 | 391 | 169 | 197 | 104 | 265 | 152 | 289 | 130 | 53 | 42 | 117 | 79 | 341 | 168 | 20 | 17 |
| Synonymous | 346 | 172 | 177 | 86 | 82 | 43 | 129 | 80 | 132 | 60 | 19 | 18 | 32 | 22 | 155 | 89 | 6 | 6 |
| Nonsynonymous | 272 | 152 | 148 | 83 | 71 | 52 | 82 | 64 | 92 | 59 | 10 | 9 | 48 | 36 | 119 | 75 | 13 | 13 |
| Stopgain | 11 | 10 | 4 | 4 | 2 | 2 | 3 | 3 | 8 | 8 | 0 | 0 | 3 | 3 | 3 | 3 | 0 | 0 |
| Stoploss | 1 | 1 | 1 | 1 | 1 | 1 | 0 | 0 | 0 | 0 | 0 | 0 | 0 | 0 | 1 | 1 | 1 | 1 |

**Figure 2.**
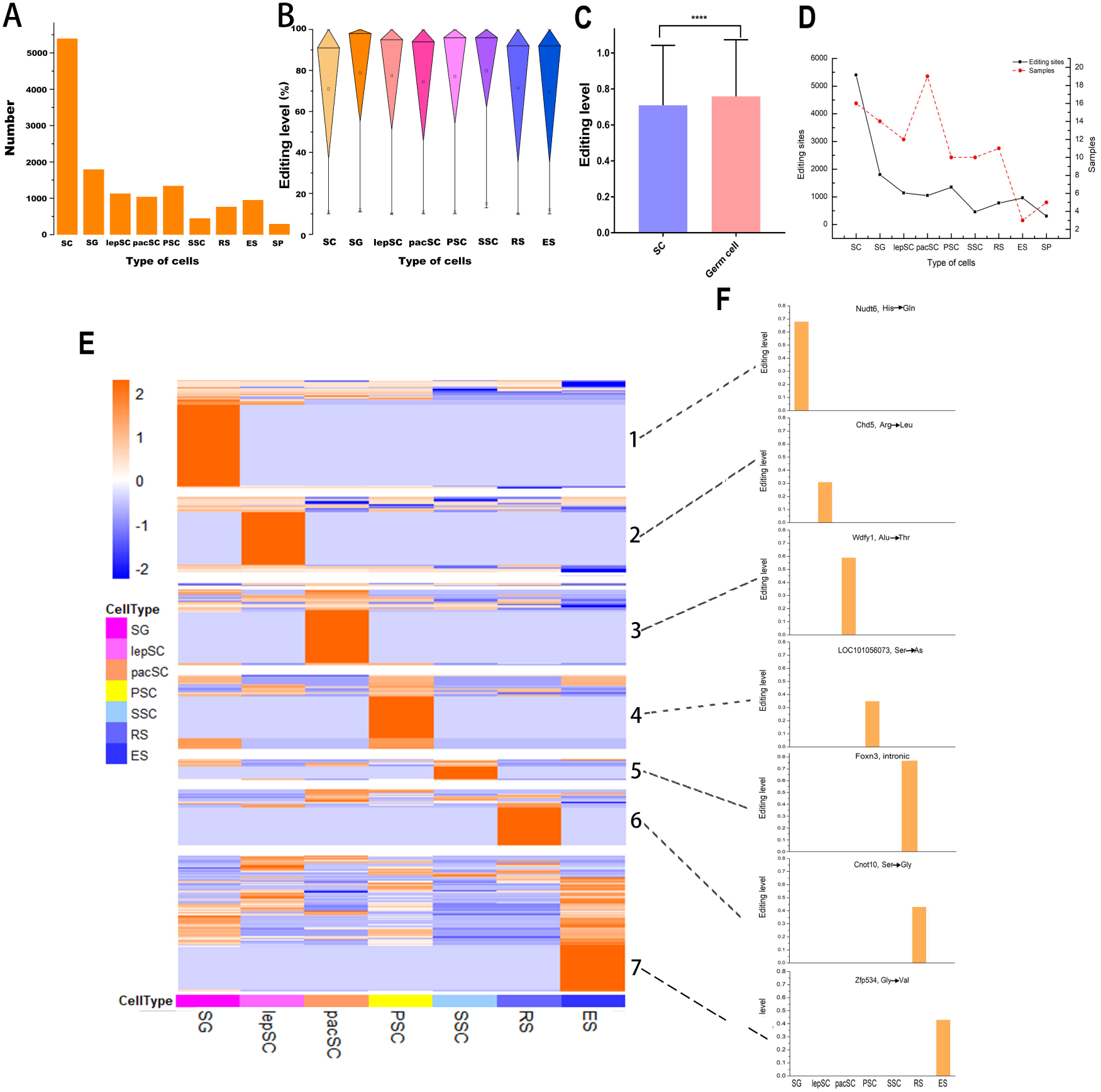
Editing sites identified in different types of cells during spermatogenesis. (A) The number of editing sites for specific types of cells. (B) Distribution of RNA editing levels in different germ cells.(C) Comparison of RNA editing levels between sertoli cells and male germ cells. Statistical signifcance was determined using *t*-test, \*\*\*\**p*< 0.0001 (D) The relationship between sample size and REs number. (E) Modular RNA editing sites based on cell type. (F) Editing levels of 7 genes that only undergo RNA editing in specific germ cells.

### Four most common types of nucleotide substitutions

Unlike humans, there are four dominant types of REs in mouse, including adenosine-to-inosine (A-to-I), guanosine-to-adenosine (G-to-A), cytidine-to-uridine (C-to-U) and uridine-to-cytidine (U-to-C)(Fig. 3A; Fig.3B; Supplemental Fig. S2). However, comparing the editing levels of these four major RNA editing events, we found that in addition to SSCs, the editing levels of A-to-I and T-to-C were higher than the other two in the other germ cells (Fig. 3C;Fig. 3D).A-to-I substitutions are significantly less abundant in mouse than in humans, mainly owing to the low representation of *Alu* repeats in its genomes(Eisenberg et al., 2005; Kim et al., 2004). Two prerequisites for A-to-I RNA editing are needed, the existence of double-stranded RNAs (dsRNAs) and Adenosine Deaminase Acting on RNA (ADAR) family of enzymes. A-to-I editing is catalyzed by members of the ADAR family of enzymes which play a role in dsRNAs (Bass, 2002)and *Alu* sequence contains long dsRNA stem loops(Sinnett et al., 1991). *Alu* sequence is the most common SINEs in humans, which is a kind of primate-specific repetitive sequences, accounts for greater than 10% of the human genome.

**Figure 3.**
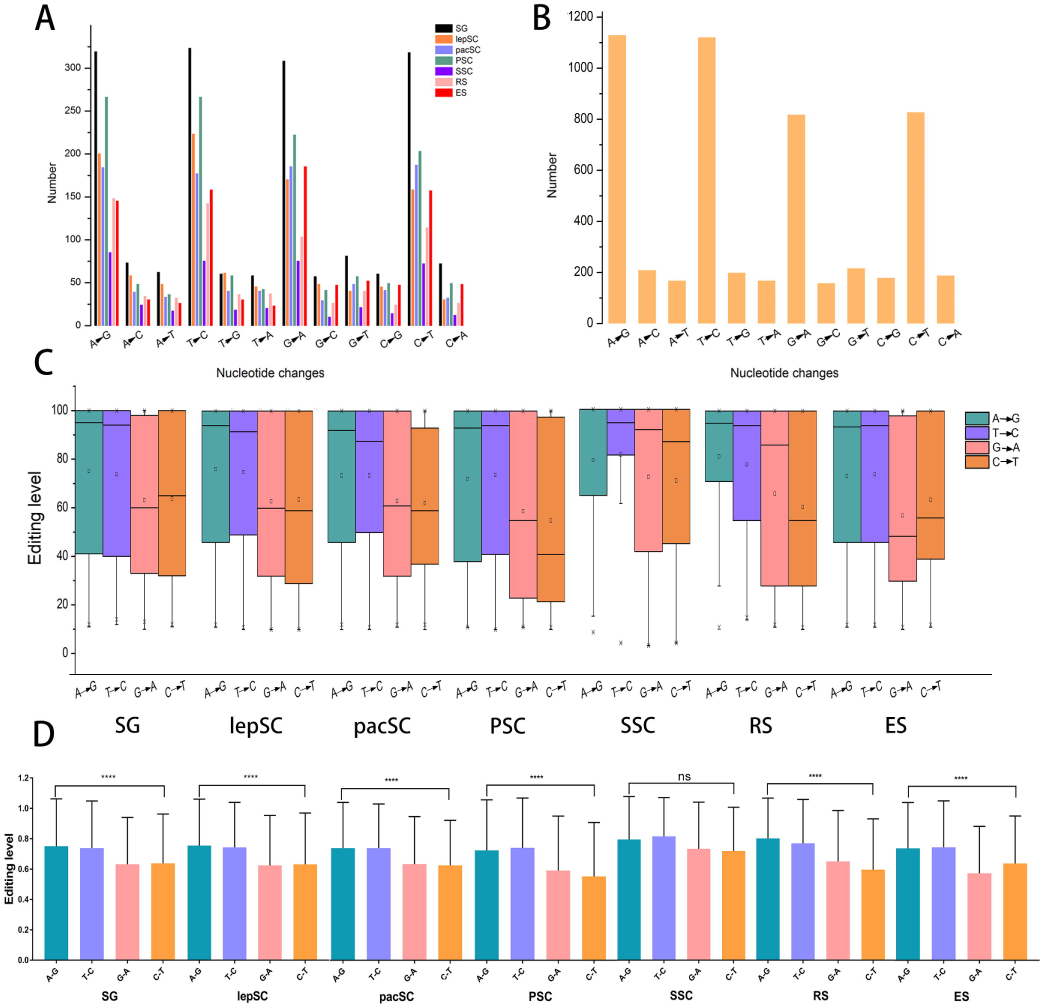
The number of 12 kinds of nucleotide changes. (A) Nucleotide changes in germ cells. (B) Nucleotide changes in sertoli cells. (C) Distribution of editing levels of four main types RNA editing in different germ cells. (D) Comparison of editing levels of these four main RNA editing. Statistical signifcance was determined usingANOVA, ****p < 0.0001, ns: non-significant.

There is no *Alu* repeats in mouse genome, but four distinct SINEs are active in mouse, including B1, B2, ID and B4. These four families in mouse occupy a smaller portion of the genome than *Alu* in human (7.6% and 10.7%, respectively). We counted the editing sites in the repetitive regions of these eight types of cells and found that REs in repetitive regions accounted for a lower proportion than in non-repetitive regions. What’s more, less editing events occur in repetitive regions in male germ cells than in sertoli cells (Table 2).

**Table 2.** Editing sites in repetitive regions

|  | SC | SG | lepSC | pacSC | PSC | SSC | RS | ES |
| --- | --- | --- | --- | --- | --- | --- | --- | --- |
| All sites | 5402 | 1803 | 1140 | 1047 | 1349 | 455 | 775 | 961 |
| Sites at repeat regions | 1122 | 196 | 179 | 139 | 107 | 49 | 72 | 67 |
| Percent | 20.7% | 10.87% | 15.7% | 13.28% | 7.93% | 10.77% | 9.29% | 6.97% |

### Editing sites ondifferent chromosomes

We analyzedthe distribution of the editing sites on all the 21 chromosomes in different types of cells (Fig.4A). Interesting, we found that editing sites showed high density on chromosome 17, the number of editing sites on per unit length (1MB) is significantly higher than other chromosomes(Fig.4B), and the same chromosomal preference can also be seen in the validation data (Supplemental Fig.S4). Functional enrichment analysis reveals that edited genes on chromosome 17 are almost immune-related (Fig.4C). As well known, take the blood testis barrier as an example, immunity play specific key roles in reproduction. The seminiferous tubules are divided into two parts by the blood testis barrier: basal and apical compartments. Meiosis, spermiogenesis and spermiation take place in the apical compartment; whereas, SG division and differentiate to pacSC occur in the basal compartment(Russell, 1977). The blood-testis barrier protects spermatocytes, sperm cells, sperm against immune system attacks and pathogenic microbes, and thus acts as an immune barrier to provide a fairly stable microenvironment(Cheng and Mruk, 2012; Jiang et al., 2014).

Compared with other chromosomes, there were a small number of editing sites on the Y chromosome and they only distributed at both ends of the Y chromosome. The validation samples also showed the same results. (Fig. 4A; Supplemental Fig. S3; Fig. 4B). Among Y chromosome genes, we found that non-synonymous editing occurs in gene *Rbmy* and *Erdr1*, and *Rbmy* has been found to be specifically expressed in adult testis (Fig. 4D). It’s reported that male mice deficient in *Rbmy* are sterile and knockout mutants of *Rbmy* are associated with azoospermia or severe oligospermia(Mahadevaiah et al., 1998; Tyler-Smith, 2008; Yan et al., 2017).

**Figure 4.**
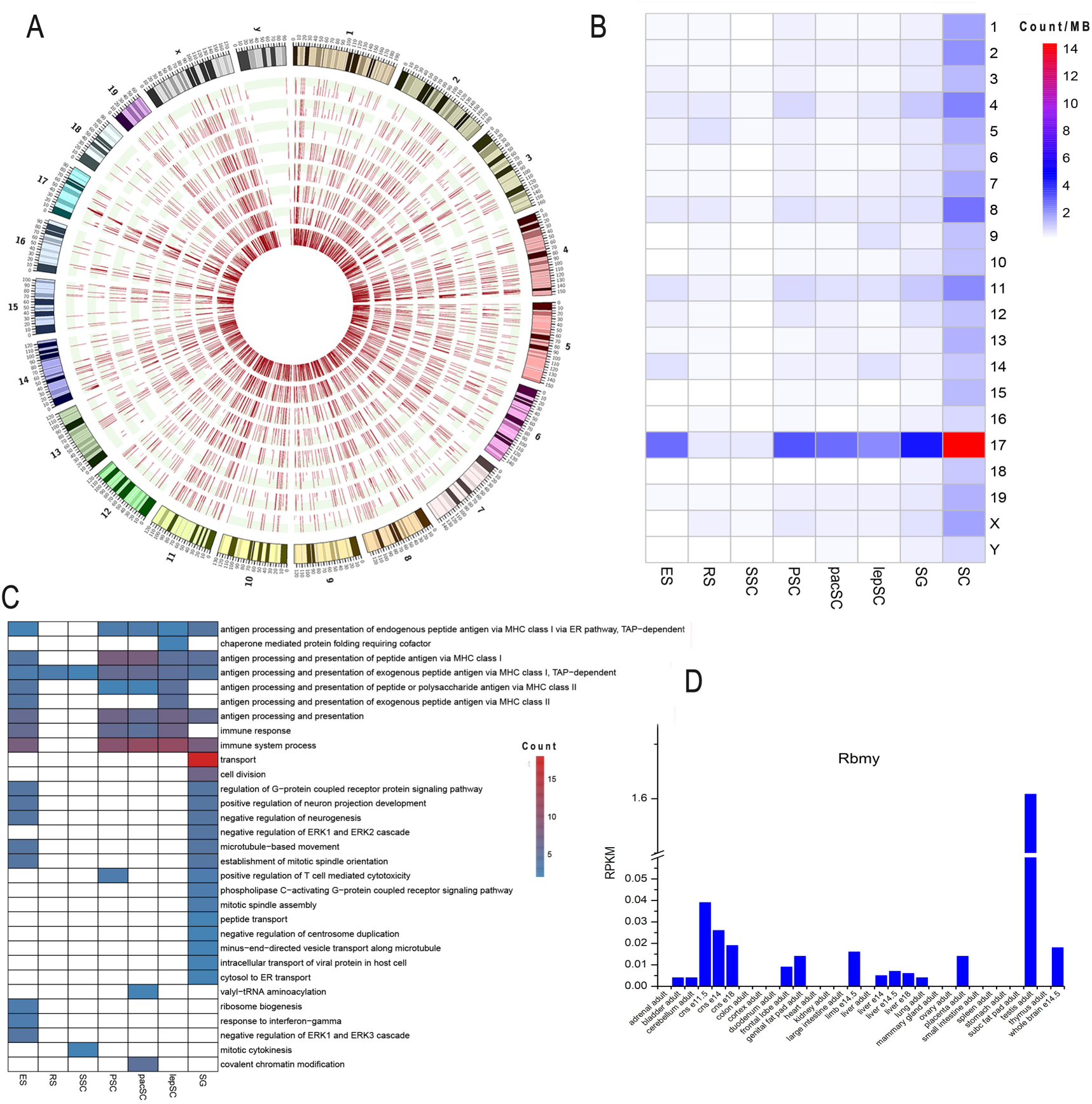
Editing sites on chromosome 17. (A) Distribution of editing sites on each chromosome in all types of cells and RNA editing levels in red bars. Cells are shown in concentric circles and ordered as follow from the inside: SC, SG, lepSC, pacSC, PSC, SSC, RS and ES. (B) The number of editing sites on the unit chromosome length of 21 chromsomes in each type of cell. (C) Function of editied genes on chromosome 17. (D) The expression of Rbmy in differenttissues.

### Half of the edited genes contain at least two editing sites

In total, REs were detected at 2012 protein coding genes in these germ cells and 2008 in sertoli cells. In male germ cells, 1042 of 2012 (69.68%) genes harbor two or more editing sites (Fig 5A; Fig 5D), while 78 genes showed more than 20 edited sites (Fig 5D). However, in sertoli cells, 998 of 2008 (49.70%) genes harbor at least two editing sites (Fig 5B). Further analysis indicates that there are more than once non-synonymous editing events at most edited genes in male germ cells (Fig 5C). There are 19 genes that contain more than 10 non-synonymous editing sites. (Fig 5E). Among them, *Gm5458*and *Gm21119*show most abundant editing events,with 77 and 70 editing sites, respectively. Interesting, we found these two genes don’t express in other tissues, but only show high expression level in adult mice testes (Fig 5F; Fig 5G), which provides strong hints that they may play key rolesduring spermatogenesis.

**Figure 5.**
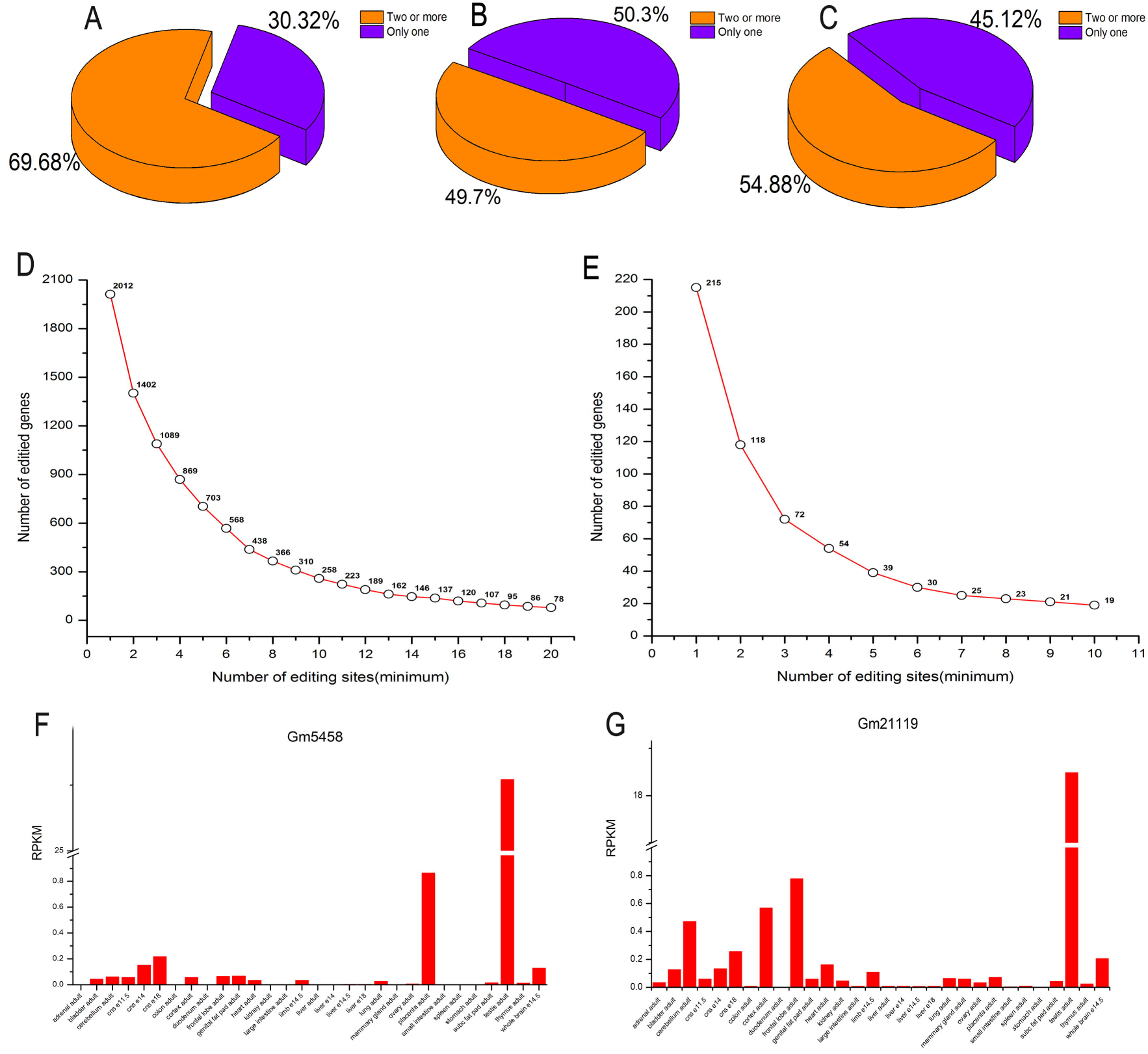
Genes with differnet numbers of editing events. (A) Proportion ofgenes with one editing event and genes with two or more editing evnets in germ cells. (B) Proportion of genes with one editing event and genes with two or more editing events in sertoli cells. (C) Proportion of genes with one nonsynonymous editing event and genes with two or more nonsynonymous editing events in germ cells. (D) The number of genes with differnet numbers of editing events in germ cells. (E) The number of genes with differnet numbers of nonsynonymous editing events in germ cells. (F)The expression of *Gm5458*across different tissues. (G) The expression of *Gm21119*across different tissues.

### Half of the editing events in CDSs lead to amino acid changes

In humans, REs primarily occurs in introns and UTRs(Bahn et al., 2012; Park et al., 2012; Sakurai et al., 2014). When analyzed the distribution of REs in different cells during spermatogenesis,we found that the editing events mainly occurred in introns, CDSs and intergenic in mouse testicular cells(Fig 6A). Further analyzed the editing sites in CDSs, to our surprise, about half of the editing events in CDSs resulted in amino acid changes (Fig 6B). On average, editing events occur in the CDSs accounted for 14%~39%. But the situation on chromosome 17 is different, on which the density of the editing sies is the highest, corresponding to 43%~62%in any germ cells, while just 33% in sertoli cells (Fig 6A; Fig 6C). Unlike sertoli cells,in male germ cells, editing events occur morefrequently in CDSs than in other regionsof chromosome 17 (Fig 6C). Chromosome 17 may be an active chromosome that performs some unclear important functions in spermatogenesis.

**Figure 6.**
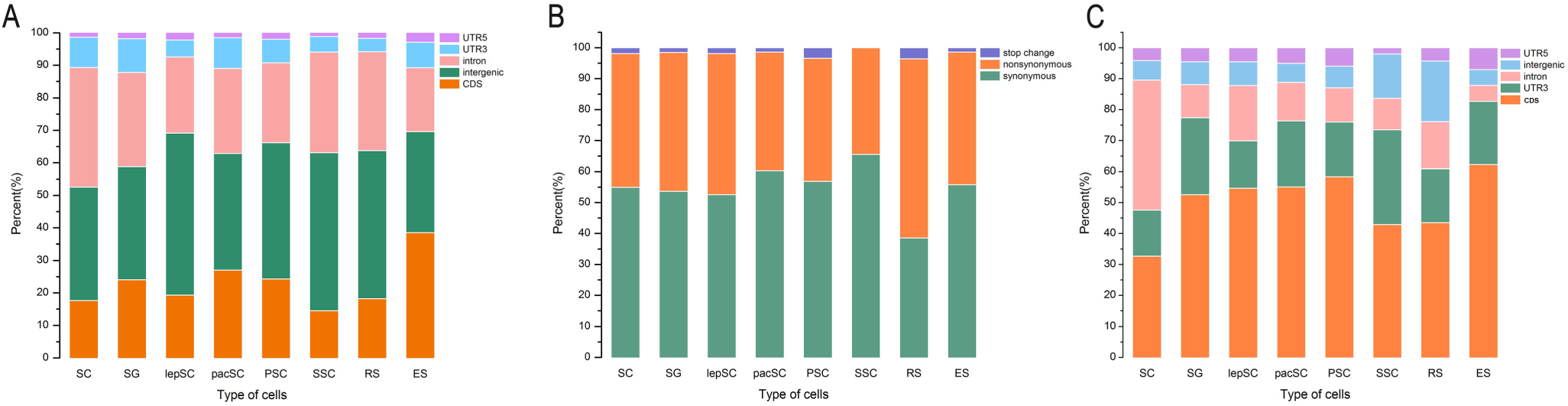
About half of the editing evnets in CDSs result in amino acid changes. (A) Distribution of the editing sites. (B) Fuctional consequences of the editing sites. (C) Distribution of the editing sites on chromosome 17.

### Function of RNA editing on spermatogenesis

We got different REs between male germ cells and sertoli cells (Fig 7A) and analyzed their related gene functions. Functional enrichment analysis results showed that many edited genes are associated with spermatogenesis, spermatid development, mitosis, cell differentiation and immunity(Fig 7B). Meanwhile, many genes have undergone REs at multiple stages of spermatogenesis, so we tried to link the relationships of these frequently edited genes with their function in spermatogenesis. Finally, many significant associations of gene-function were detected among all the seven developmental stages, and we showed some typical gene-function pairs here (Fig 7C). Both *Ddx3x* and *Hjurp* are found to be related to chromosome segregation. *Ddx3x* has undergone REs in the seven stages of spermatogenesis, whereas *Hjurp* has been edited in four stages. In addition, *Adad1* and *Rnf17* are involved in spermatogenesis and spermatid development, while *Cxcr4* and *Cxcl12* play roles in germ cell migration. These four genes are also edited in different stages of spermatogenesis. In short, these edited genes show close relationships with spermatogenesis through REs.

**Figure 7.**
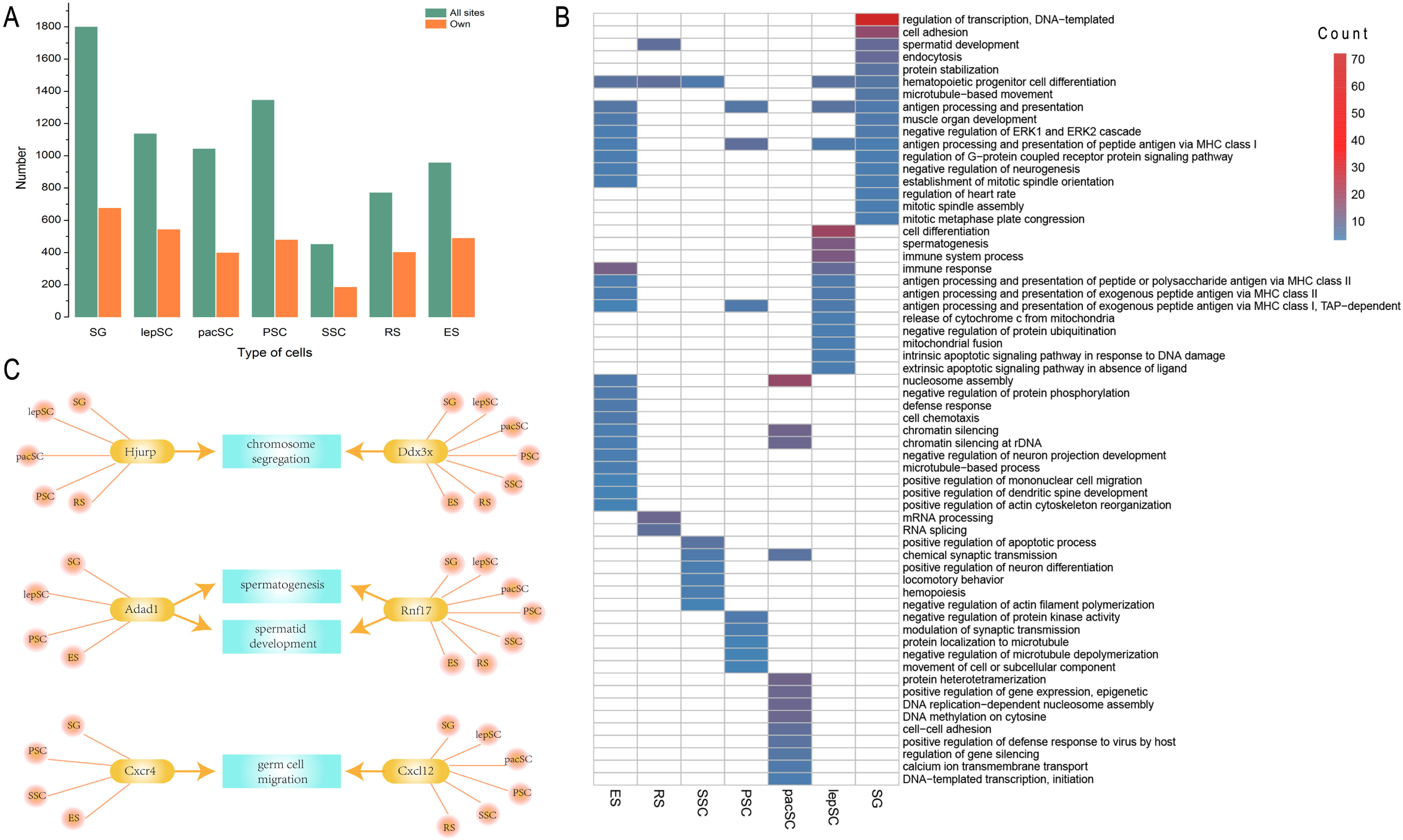
Significant editing events for spermatogenesis. (A) All sites in male germ cells and eiting sites only in germ cells. (B) Function of genes that have editing sites only in germ cells. (C) The function of genes which are edited in multiple stages.

### Validation of conserved RNA editing site between species and tissues

Interesting, we found that there is a non-synonymous A-to-I RNA editing site in *Cog3* during mouse spermatogenesis, which have been reported previously in brains of human, rhesus and mouse(Li et al., 2009; Ramaswami and Jin, 2014). This result demonstrated that the editing site is not only conserved between species, but also conserved across tissues. To validate the authenticity of our results, we sequenced the genome and cDNA of *Cog3* in male germ cells. By comparing the nucleotides of the genomeand cDNAat this site(Fig 8), we found that, as we detected earlier based on integrated RNA-Seq, the A-to-I RNA editing of *Cog3*, still occurs during mouse spermatogenesis which indicates an important role for the development of male germ cells.

**Figure 8.**
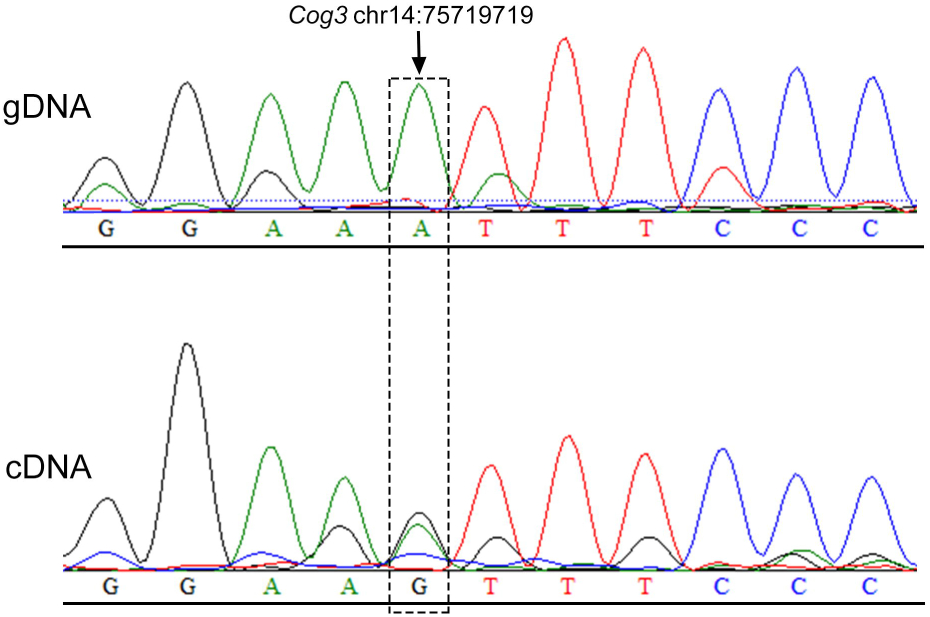
Validation of RNA editing site withSanger sequencing. Sanger sequencing of cDNA and gDNA, at the site chr14:75719719 of gene *Cog3*the genomic A is highly edited to I.

## DISCUSSION

In this study, we integrated all the published RNA-Seq data of all kindes of male germline cells with sertoli cells as control during mouse spermatogenesis. Based on the power of integrated data, we established the robust landscape of REs during the development of spermatogenesis. Finally, 7530 editing sites were identified in total, 2012 genes in male germ cells have presented editing events, and more than half of them harbor at least two editing sites. The two most abundant editing genesare *Gm5458* and *Gm21119*, with 77 and 70 editing sites, respectively. Both of themwere foundspecificallyexpressed in testis.

Unlike human, A-to-I substitutions does not showan absolute predominantein mouse male germ cells(Levanon et al., 2005), and there are four most common types of nucleotide substitutions, they are A-to-I, G-to-A, C-to-T and T-to-C. Editing sites are enriched on some regions of chromosome, with chromosome 17 as the highest density chromosome. On the Y chromosome, editing events occur specifically at both ends of the chromosome, and non-synonymous editing events only occur on the genes *Rbmy* and *Erdr1*, where *Rbmy* plays an important role in sperm formation. We hypothesize that *Rbmy* plays a role in spermatogenesis through REs as a post-transcriptional regulator. On the whole, unlike human,REs mainly occurs in introns, intergenic and CDSs in mose,and about half of the editing events in CDSs can lead to amino acid changes. As to chromosome 17, enrichmenst of editing events in CDSs were most obvious. Intereting, the editied genes on chromosome 17 are almost immune-related, while as well known, immunity and reproductive are very closely linked.Therefore, we have reason to speculate that chromosome 17 may be a key chromosome during spermatogenesis through immunie systems.What’s more, edited genes are found to be related to spermatogenesis, spermatid development, mitosis, cell differentiation and immunity. Among them,some are frequently edited at multiple stages,such as *Ddx3x*, *Hjurp*, *Adad1*, *Rnf17*, *Cxcr4* and *Cxcl12*. Summary, we integrated all published RNA-Seq datasets and established the landscape of REs during spermatogenesis. Based on the landscape, we desconstructed molecular mechanisms for the development of male germ cells through REs from the perspective of post-transcriptional regulation.

In order to take advantages of the power of merging different studies as one union, we integrated all the published RNA-Seq datasets at the same conditions. As a typical dynamic complex process, spermatogenesis was chose to apply our strateges. Our results showed that there are much novel insights can be detected by integrating different datasets. Our strateges should be general for other researches to make use of the huge mount of published biological datasets. Beyond RNA-Seq data, our strateges can be also used for other types of sequencing datasets.

This work focused on the REs regulation for mRNA of coding proteins. But as well known, the majority of RNAs generated in the cell are non-coding RNAs (ncRNAs). In human, it is estimated that about two-third of genome is transcribed and only 2% of these transcribed RNAs are translated into proteins. The functions of ncRNAs vary from regulating transcription, translation, DNA repair and cell fate decision(Bunch, 2017).It’s recently reported that non-coding RNA may be associated with cytoplasmic male sterility in plant(Stone et al., 2017). What roles do non-coding RNAs play in spermatogenesis and whether they affect the formation of spermatozoa by REs are still unclear. These problems should be considered in the future researches and it will be promising for the molecular mechanism exploring of reproduction.

## MATERIALS AND METHODS

### Data collection and classification integration

RNA-seq reads of mouse spermatogenesis used to identify editing sites were downloaded from GEO and SRA, and then categorized them into nine groups according to their development stages, including spermatogonia, leptotene spermatocytes, pachytene spermatocytes, primary spermatocytes before meiosis □, secondary spermatocytes, round spermatids, elongative spermatids, sperm and sertoli cells as control. Validation datasets of spermatogonia and round spermatids was under accession number GSE75826.

### Read mapping and identification of RNA editing sites

RNA-seq reads were aligned to the complete genome of mouse available in Ensembl with program HISAT2(Kim et al., 2015). RNA editing sites were identified by REDItools(Picardi and Pesole, 2013)and a series of stringent filter were implemented to eliminate false positive, as was described in Supplemental Methods. Known variations were downloaded from UCSC and were moved by bedtools, with details in Supplemental Methods.

### Annotation of RNA editing

Annovar(Wang et al., 2010)was used to annotate RNA editing based on their related gene, while the related annotation files refGene.txt and chromFa.tar.gz were downloaded from UCSC with version mm10. Based on these data, gene-based transcriptional expression and the distribution of editing sites on chromosomes, genes and genomic elements were caculated.

### Calculation of gene expression

After mapping with HISAT2, the alignments files were passed to StringTie for transcript assembly and calculating the expression level of transcript. Finally, Ballgown took all the transcripts and abundances from StringTie and counted expression based on genes(Pertea et al., 2016), some details in Supplemental Methods.

### Mice

Five adult male Kunming mice were used in our experiments. All mice were housed in a barrier facility under natural light at 20-25 ℃, free access to food and water. All experimental procedures involving animals were approved by the Institutional Animal Care and Use Committee of Northwest A&F University.

### Isolation of germ cells

The convoluted tubule was isolated from mouse testicles in a sterile environment and digested with trypsin to obtain single cell viability. Cells were cultured using DMEM/high glucose (Gibco, Grand Island, NY, USA) and supplemented with 10% FBS (Gibco), 100 U/ml penicillin, 100 mg/ml streptomycin, 1 MEM non-essential amino acids. After 4 hours, when sertoli cells adhered to the bottom, germ cells were collected by centrifugation(500g/5min).

### PCR and Sanger sequencing analysis

The genome was obtained through TIANamp Genomic DNA Kit and the total RNA was extracted by RNAiso Plus using a one-step method of isothiocyanate guanium-phenol-chloroform. PCR amplification was performed using the cDNA and genome.

The primer sequences were as follows: DNA primer F: ACTGCAGAGAACTATCACTTTGAGG, R: AAGATGATGTCACTCTGTTCCCTTT, cDNA primer F: TAGATGCATAGATAGGGCGGTGT, R: TGCTGTGAGAGGGTGTACTTGG. The PCR products were sequenced by Sanger sequencing analysis in Sangon biotech.

